# Spatiotemporal Transcriptomic Dissection Uncovers Age-Dependent Deceleration of Esophageal Cell Differentiation

**DOI:** 10.64898/2026.05.18.726035

**Authors:** Jinho Jang, Jie Zhang, Amy S. Park, Sohee Jun, Jae-Il Park, Kyung-Pil Ko

**Affiliations:** Department of Experimental Radiation Oncology, Division of Radiation Oncology, The University of Texas MD Anderson Cancer Center, Houston, TX 77030, USA; Graduate School of Biomedical Sciences, The University of Texas MD Anderson Cancer Center, Houston, TX 77030, USA; Program in Genetics and Epigenetics, The University of Texas MD Anderson Cancer Center, Houston, TX 77030, USA

**Keywords:** esophagus, stem cells, aging, single-cell RNA-sequencing, spatial transcriptomics

## Abstract

Precise orchestration of stem and progenitor cells is essential for tissue homeostasis and regeneration but becomes dysregulated during aging. Despite the known markers, the age-related dynamics of esophageal epithelial cell lineages remain unclear. Using single-cell single cell transcriptomics, we analyzed human esophageal epithelia across different age groups. We identified two stem cell populations: quiescent (qeSCs) and proliferative (peSCs) esophageal stem cells. qeSCs from young donors showed higher *WNT10A* expression and Wnt signaling activity. Analysis of cell lineage trajectories combined with cell plastic potentials showed stronger connectivity between peSCs and differentiated cells in younger tissues, indicating more efficient and rapid epithelial turnover and homeostatic maintenance. Cell-cell interaction analysis further demonstrated that NOTCH signaling is more prominent within peSCs and qeSCs in younger esophagi, whereas in older tissues, NOTCH activity is preferentially retained in differentiated cells. Additionally, the inflammatory signaling, Interleukin-1 pathway, is more active in younger esophagi but is largely restricted to differentiated cells. Our findings suggest that age-related decline in esophageal homeostasis is primarily driven by impaired differentiation dynamics rather than by alterations in stem cell self-renewal capacity.

## Introduction

The esophagus is a tubular organ that transports food and fluids to the stomach through coordinated peristaltic movement. The esophageal epithelium consists of multiple layers of cells, with basal cells located at the bottom layer where stem cells reside.^1^ Proliferating cells in the basal cell layer generate daughter cells that migrate toward the lumen and progressively differentiate into mature squamous epithelial cells.^2,3^ To maintain epithelial integrity and function, esophageal epithelial cells undergo continuous turnover approximately every 11 days in human, requiring tightly regulated stem cell activity for normal tissue homeostasis.^3^ However, this homeostatic balance is disrupted during tumorigenesis, where cancer stem cells gain clonal dominance and drive malignant progression.^4-6^ Therefore, understanding normal esophageal stem cell biology and lineage trajectories is essential for distinguishing normal from cancer stem cells and for developing strategies to prevent tumor initiation. Notably, esophageal cancer is strongly associated with aging,^7^ highlighting the importance of investigating age-related changes in epithelial cell lineage dynamics.

Previous studies have identified normal esophageal epithelial stem cells based on marker gene expression, including SOX2, TRP63, KRT15, ITGA6, ITGB4, and CD73.^2,8,9^ However, the expression of these markers is broadly distributed across basal epithelial layers, limiting their specificity and practical utility for definitive stem cell identification.

Advances in single-cell RNA sequencing (scRNA-seq) technologies have enabled more refined characterization of epithelial stem cell populations, leading to the identification of quiescent basal cells that are thought to serve as a reservoir for proliferating progenitor cells. Markers such as COL17A1, DST, and PDPN have been proposed to identify these populations.^10-12^ However, many of these studies relied on tissue adjacent to diseased regions as normal controls, which may complicate interpretation. Emerging evidence suggests that histologically normal tissues adjacent to lesions can exhibit distinct transcriptomic alterations compared with healthy normal tissues.^13^ Furthermore, age-dependent characteristics of esophageal epithelial stem cells remain insufficiently characterized.

In this study, we performed single-cell transcriptomic analyses of healthy normal esophageal epithelial cells and compared their stem cell properties and lineage trajectories across different age groups, with experimental validation in murine tissues by immunostaining and spatial transcriptomics.

## Results

### Limitations of conventional esophageal stem cell markers in mapping esophageal cell lineages

To investigate the cellular origins of normal esophageal epithelium, we integrated single-cell RNA-seq datasets from eight healthy donors derived from two independent studies^1,12^. After isolating epithelial cells, datasets were integrated and batch-corrected using the Harmony algorithm. Epithelial cells were clustered by unbiased Leiden algorithm and tested with known marker genes of differentiated cells and stem cell marker genes of esophagus (**Fig. 1A–C**). Interestingly, previously reported esophageal stem cell (eSC) marker genes were not restricted to a discrete epithelial cluster. Instead, these markers were broadly expressed across multiple epithelial clusters and showed substantial overlap with differentiated cell markers, suggesting limited specificity as stem cell identifiers. Conventional eSC markers, including SOX2 and TP63, exhibited widespread expression throughout the epithelial compartment.

**Figure 1.**
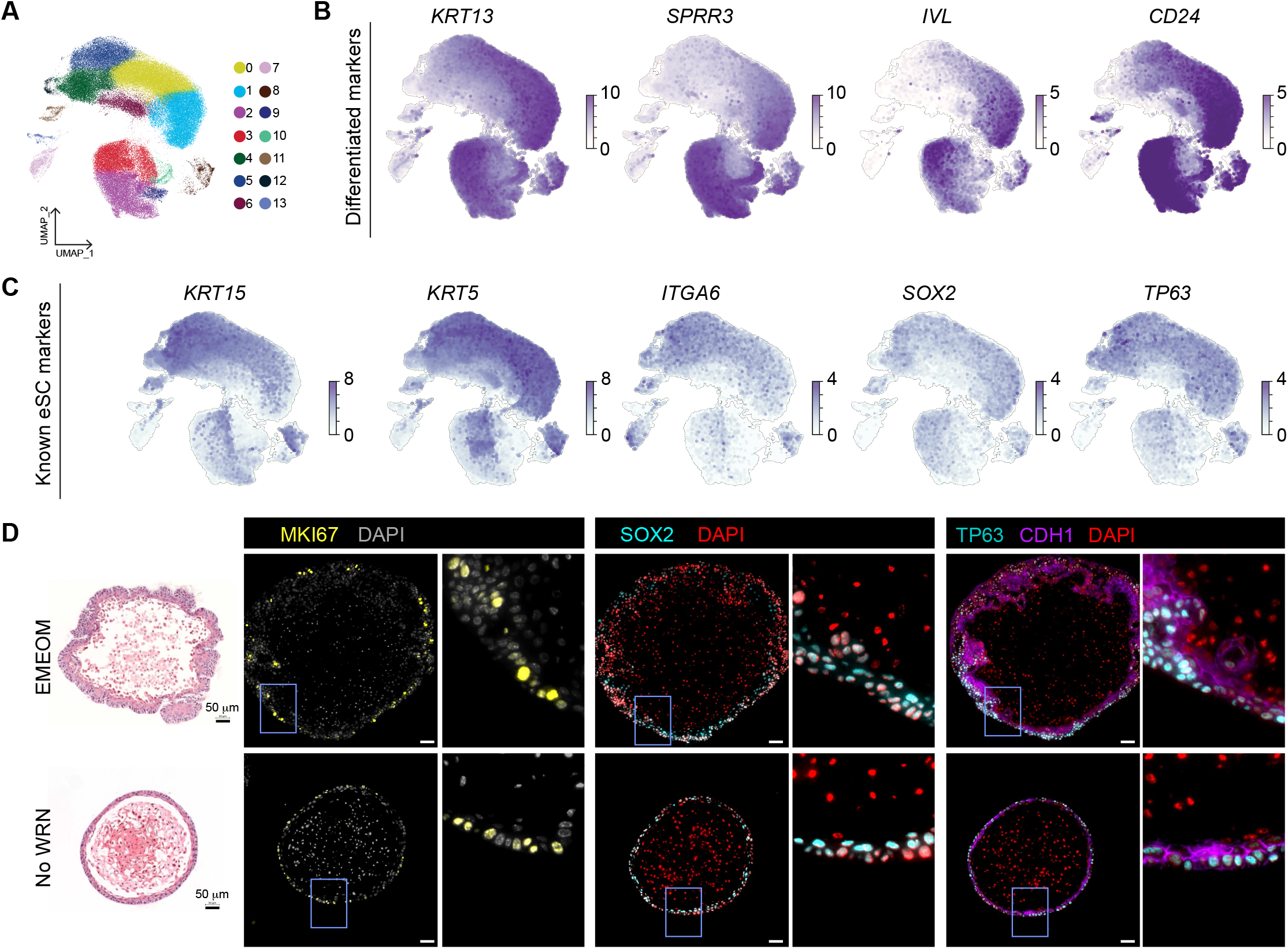
Stem cell markers in normal human esophageal epithelium. **A**. UMAP visualization of the integrated single-cell RNA-sequencing dataset generated from esophageal epithelial tissues of eight healthy donors. Following dataset integration, epithelial cell populations were extracted and analyzed. **B**. UMAPs showing the expression of differentiated epithelial cell marker genes. **C**. UMAPs displaying the expression of previously reported esophageal epithelial stem cell markers. **D**. Representative H&E images (left) and immunohistochemical staining for MKI67, SOX2, TP63, and CDH1 in murine esophageal organoids cultured under stemness-enriched conditions (EMEOM) or homeostatic conditions without WRN supplementation (No WRN). Scale bars, 50 µm.

To examine the spatial distribution of these markers, we performed immunofluorescent staining in esophageal organoids (EOs). Consistent with the transcriptomic findings, SOX2 and TP63 expression was not confined to a subset of cells and extended beyond proliferative cells represented by MKI67 (**Fig. 1D**). Rather, both markers were broadly expressed across the basal cell layer. To determine whether this pattern resulted from enhanced stem cell activity driven by organoid culture medium (EMEOM), we removed Wnt3a, R-spondin 3, and Noggin (WRN) from EMEOM to better mimic homeostatic esophageal tissue. Although WRN deprivation led to thinning of the basal layer compared to EMEOM conditions, SOX2 and TP63 expression remained broadly distributed within basal cells (**Fig. 1D**). These findings highlight the limited specificity of conventional eSC markers for defining the cell of origin in normal esophageal epithelium.

### Molecular signatures of quiescent and proliferative stem cells in the aging esophagus

To infer the cells of origin within normal esophageal epithelium, we applied RNA velocity-based lineage trajectory analysis to the integrated single-cell transcriptomic dataset (**Fig. 2A, B**). The inferred terminal states corresponded well with differentiated epithelial cells based on established marker gene expression. In contrast, the inferred root population was confined to specific epithelial clusters, suggesting distinct stem-like subpopulations. We next compared previously reported stem cell and proliferation markers with the RNA velocity-defined clusters. Proliferation-associated genes, including *MKI67, STMN1*, and *PCLAF*, were enriched in the second cluster (blue), consistent with an actively cycling progenitor-like population (**Fig. 2C**). In contrast, stem cell markers identified in prior scRNA-seq studies, such as *COL17A1, DST*, and *PDPN*, were preferentially expressed in the first cluster (green). Given that *COL17A1* and *DST* have been previously annotated as markers of quiescent basal cells, we designated the first cluster as quiescent esophageal stem cells (qeSCs) and the second cluster as proliferating esophageal stem cells (peSCs). Based on RNA velocity trajectories and marker gene expression, we classified esophageal epithelial cells into four populations: qeSCs, peSCs, intermediate epithelial cells (Intermediate epi), and differentiated epithelial cells (Differentiated epi) (**Fig. 2D**). Although *COL17A1, DST*, and *PDPN* demonstrated improved resolution compared with conventional markers such as *SOX2* and *TP63, COL17A1* and *DST* still exhibited broad expression across multiple epithelial populations, including proliferating cells.

**Figure 2.**
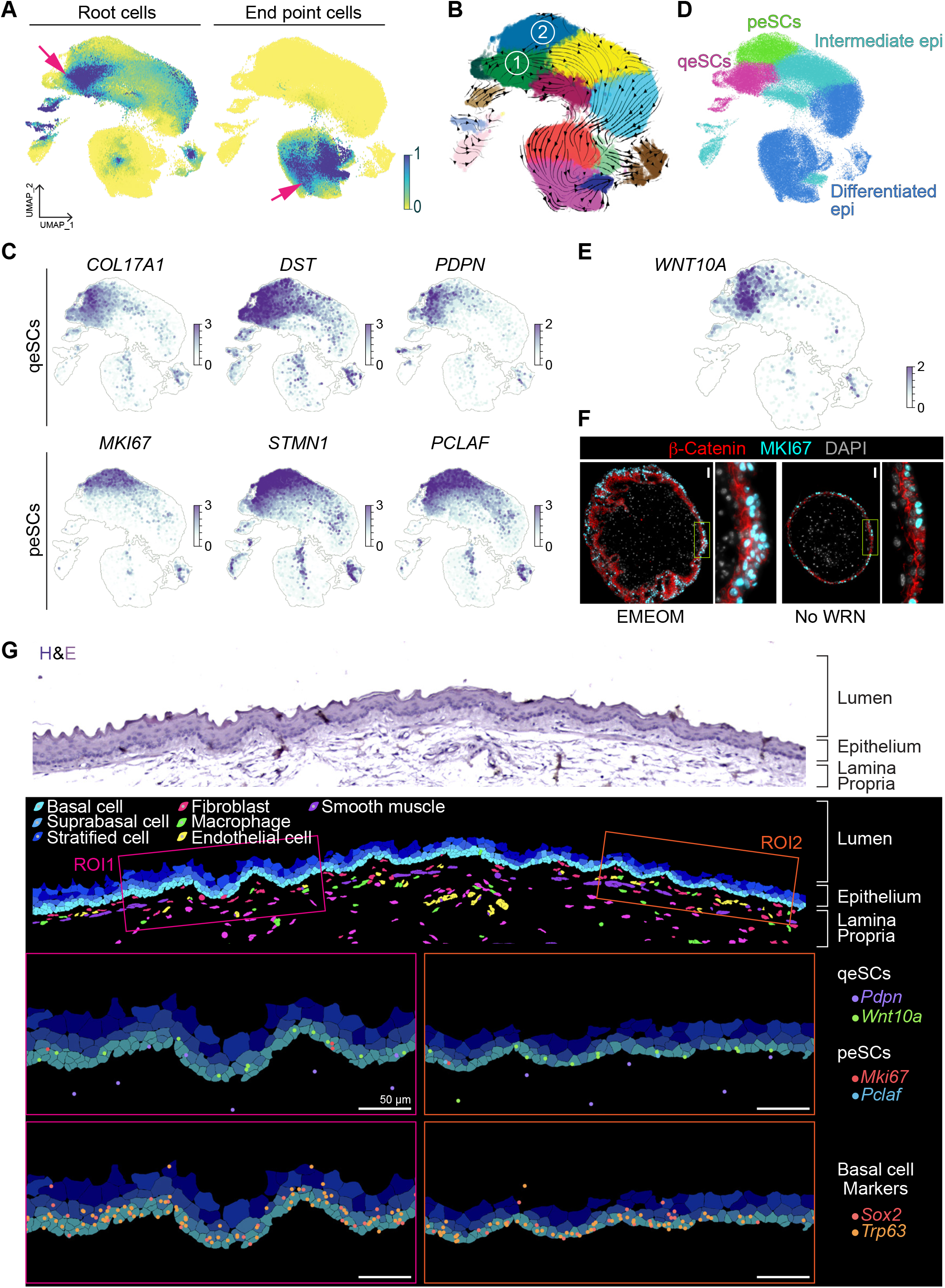
Identification of distinct stem cell populations in normal esophageal epithelium. **A**. UMAP visualization showing RNA velocity-defined root cells and differentiation endpoints. **B**. Two distinct stem cell populations identified as potential cells of origin based on RNA velocity streamlines. **C**. UMAPs displaying expression of previously reported quiescent stem cell markers (*COL17A1, DST*, and *PDPN*) and proliferating stem cell markers (*MKI67, STMN1*, and *PCLAF*). **D**. UMAP showing epithelial cell-type annotation based on marker gene expression patterns. **E**. UMAP displaying WNT10A expression. **F**. Immunohistochemical staining showing β-catenin expression in murine esophageal organoids. **G**. Representative H&E staining and spatial transcriptomics-based images of normal murine esophageal tissue. Expression patterns of established stem cell markers and *Wnt10a* are shown. Scale bars, 50µm.

To identify more specific qeSC markers, we performed differential gene expression analysis across the annotated clusters. This refined analysis identified *WNT10A* as a marker with higher specificity for qeSCs compared to previously reported genes (**Fig. 2E**). Consistently, downstream β-catenin signaling activity, assessed by nuclear β-catenin localization, was restricted to a small subset of basal cells in homeostatic esophageal organoids (**Fig. 2F**), supporting specific WNT pathway activation within the qeSC population.

To further validate these findings in vivo, we examined spatial transcriptomic data from normal mouse esophagus (**Fig. 2G**). As observed in human scRNA-seq data, *Sox2* and *Trp63* were broadly expressed across the basal layer. In contrast, *Wnt10a* exhibited confined expression within a limited subset of basal cells, consistent with a restricted qeSC population. Although *PDPN* appeared enriched in qeSCs in scRNA-seq data, its expression in spatial transcriptomics was minimal within epithelial cells and was instead more prominent in cells located in the lamina propria.

These findings identify WNT10A as a more specific marker of quiescent esophageal stem cells and support a hierarchical organization of normal esophageal epithelium defined by distinct quiescent and proliferative stem cell states.

### Age-dependent deceleration of esophageal epithelial cell differentiation

To investigate potential age-related differences in the cells of origin within esophageal tissue, we separated the datasets according to donor age groups (10–15, 30–35, 40–45, 50–55, 60–65, and 65–70 years) and compared their cell lineage trajectories. RNA velocity–based lineage analysis did not reveal substantial differences in the overall trajectory patterns, which consistently originated from qpSC or peSC populations and progressed toward intermediate and differentiated cells across all age groups (**Fig. 3A**). In addition, the proportions of each cell cluster were not significantly altered by age (**Supplementary Fig. S3A**), and similar cell-cycle patterns were observed among all age groups (**Supplementary Fig. S3B, C**). However, *WNT10A* expression within the qeSC cluster was highest in the 10–15 age group, suggesting age-dependent differences in stem cell activity or regenerative potential (**Fig. 3B**).

**Figure 3.**
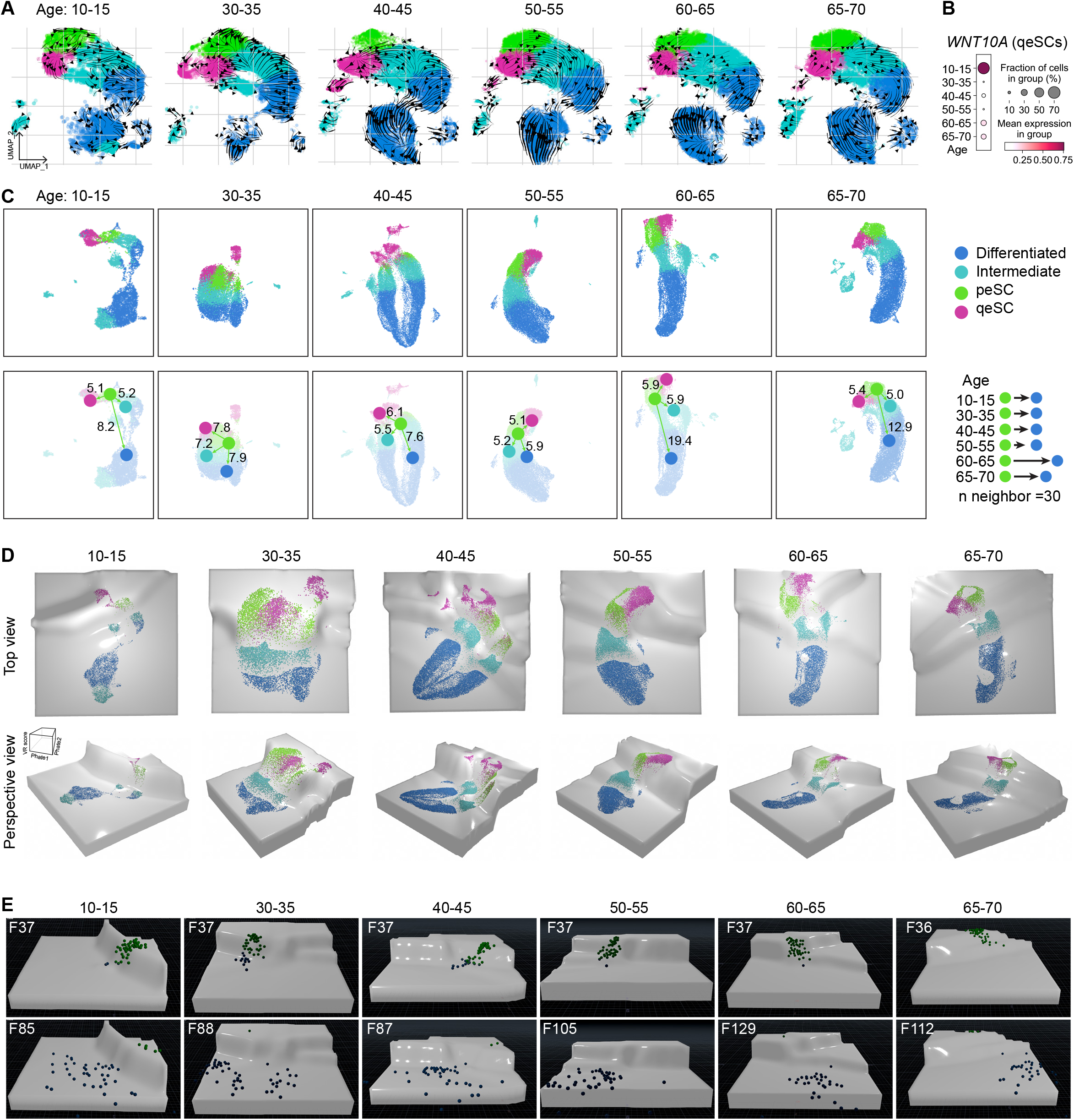
Age-dependent alterations in epithelial cell differentiation. **A**. RNA velocity–based differentiation trajectories visualized on UMAPs across different age groups. **B**. Dot plot showing *WNT10A* expression across age groups. **C**. Reconstructed UMAPs generated separately for each age group using RNA velocity analysis. Average neighboring-cell distances between cell types are displayed. **D**. Waddington’ landscape projections generated using VR scores across different age groups. **E**. Cell differentiation simulations performed on Waddington landscapes according to age group. Starting and endpoint frame numbers are indicated.

To determine whether aging influences differentiation probability, we evaluated the distances between cell clusters across age groups. Because UMAP embeddings were generated separately for each age dataset, we calculated the distances between neighboring cells of different cell types within each embedding (**Fig. 3C**). By averaging these cell-to-cell distances, we identified notable differences in tissues derived from donors aged 60–65 and 65–70 years. In these older age groups, the distance between peSCs and differentiated cell cluster was greater than that observed in younger groups, suggesting a reduced probability of differentiation from peSCs into differentiated cells.

To further model differentiation potential, we generated Waddington landscapes based on valley-ridge (VR) scores (**Fig. 3D**), as we recently performed.^14,15^ To estimate differentiation dynamics, we simulated cell-state transitions by releasing 50 balls from the ridge region corresponding to qeSC and peSC populations and measured the time required for the final ball to reach the differentiated cell state. Differentiation time was quantified in animation frames and compared across age groups. Younger groups (10–15, 30–35, and 40–45 years) required approximately 48–51 frames to complete differentiation, whereas older groups exhibited progressively prolonged differentiation times: 50–55 years required 68 frames, and the 60–65 and 65–70 groups required 76–92 frames (**Fig. 3E** and **Supplementary Video 3A–F**). These findings suggest that aging is associated with delayed differentiation dynamics in esophageal epithelial cells.

### Age-related rewiring of NOTCH and inflammatory signaling in the esophageal epithelia

To investigate signaling cues involved in normal esophageal epithelial homeostasis, we performed CellChat analyses to evaluate the cell-to-cell interactions (**Fig. 4A**). From the integrated epithelial-immune cell datasets, we identified several significant signaling pathways, including EGF, NOTCH, NRG, and IL1, that were enriched in association with peSCs and qeSCs (**Fig. 4B**). Although qeSCs and peSCs were predicted to interact with multiple cell types, including immune cells and fibroblasts, both populations showed highly consistent interaction patterns involving the NOTCH signaling pathway specifically within epithelial cell populations (**Fig. 4C**).

**Figure 4.**
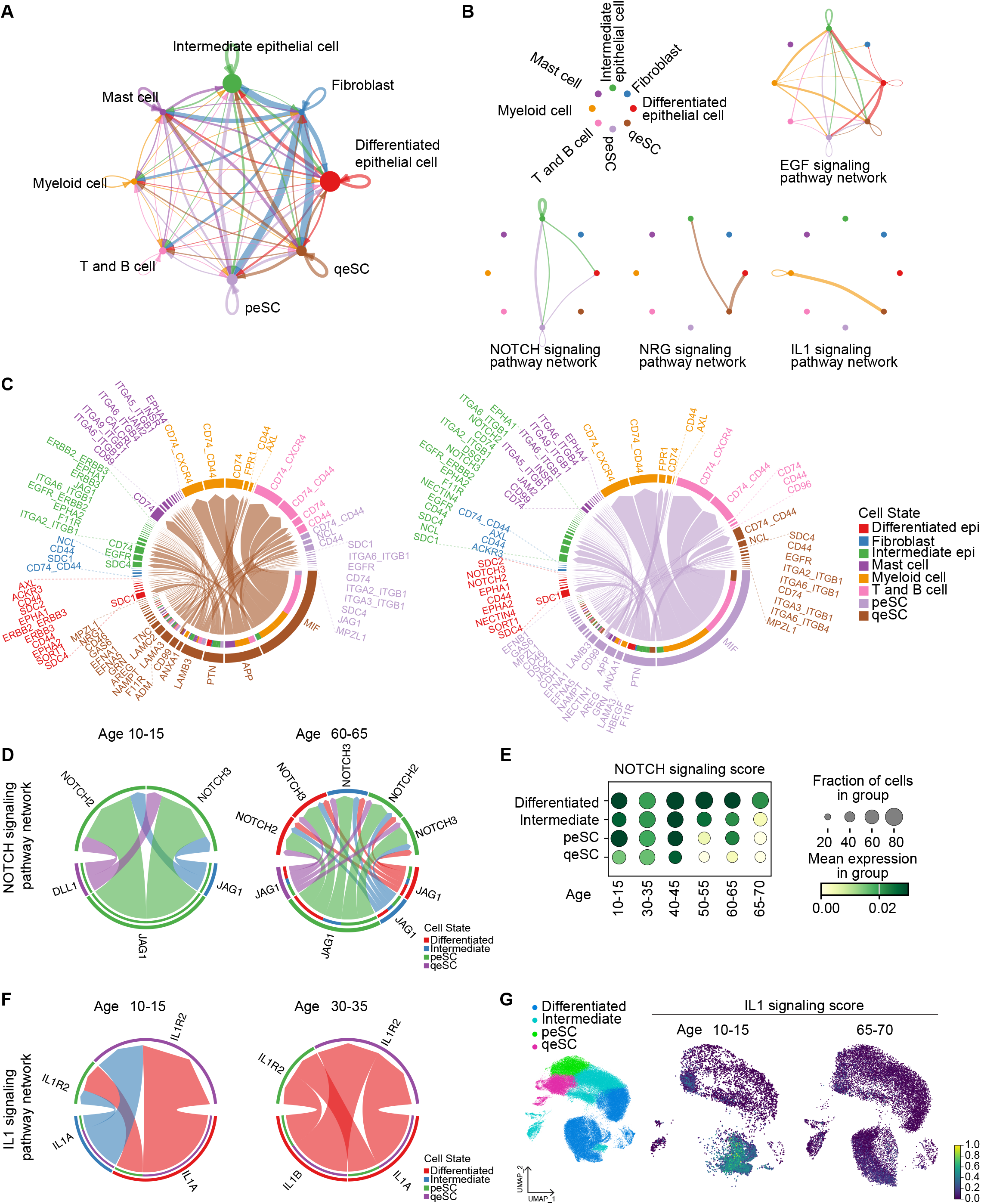
Changes in stem cell signaling pathways during esophageal aging. **A**. Cell–cell interaction networks inferred by CellChat analysis using integrated datasets containing epithelial and non-epithelial cell populations. **B**. Significant signaling pathways enriched in association with qeSCs and peSCs. **C**. Molecules involved in qeSCs- and peSCs-mediated epithelial cell–cell interactions. **D**. NOTCH signaling–mediated epithelial cell interactions in tissues from the 10-15 and 60-65 age groups. **E**. NOTCH signaling activity scores across epithelial cell populations and age groups. **F**. IL1 signaling–mediated epithelial cell interactions in the 10-15 and 30-35 age groups. **G**. IL1 signaling activity scores in epithelial cell populations from the 10-15 and 60-65 age groups.

Therefore, we examined whether NOTCH-mediated epithelial cell–cell interactions differed across age groups. Strikingly, NOTCH signaling interactions in the youngest group (10–15 years) were predominantly enriched within peSCs and qeSCs, whereas in older tissues (60– 65 years), NOTCH signaling was distributed more broadly and was increasingly mediated through differentiated epithelial cells (**Fig. 4D** and **Supplementary Fig. S4A**). These observations were further supported by NOTCH signaling activity scores within epithelial cells. Although NOTCH signaling activity was broadly detected in differentiated cells across all age groups, enrichment within peSCs and qeSCs was primarily observed in younger tissues (10–15, 30–35, and 40–45 years) and progressively declined in older groups (50–55, 60–65, and 65–70 years) (**Fig. 4E**).

Comparative cell-cell interaction analysis also revealed age-associated differences in IL1-mediated signaling. Cell-cell interactions mediated by IL1 signaling were enriched in younger tissues (10–15 and 30–35 years) but were markedly reduced in older age groups (**Supplementary Fig. S4B**). Predicted signaling patterns suggested that intermediated or differentiated cells-derived ligands such as *IL1A* and *IL1B* were received by peSCs and peSCs (**Fig. 4F**). Interestingly, these signal-receiving peSCs and qeSCs predominantly expressed *IL1R2*, a decoy receptor for IL1 signaling. These findings suggest that, although IL1 signaling activity is elevated in younger tissues, signaling output may be attenuated within peSC/qeSC populations while remaining stronger in differentiated epithelial cells (**Fig. 4G** and **Supplementary Fig. S4C**).

Together, these findings suggest that younger esophageal epithelium exhibits distinct cell-cell interaction networks, particularly involving qeSC and peSC populations, that are mediated through NOTCH and IL1 signaling pathways (**Fig. 5**).

**Figure 5.**
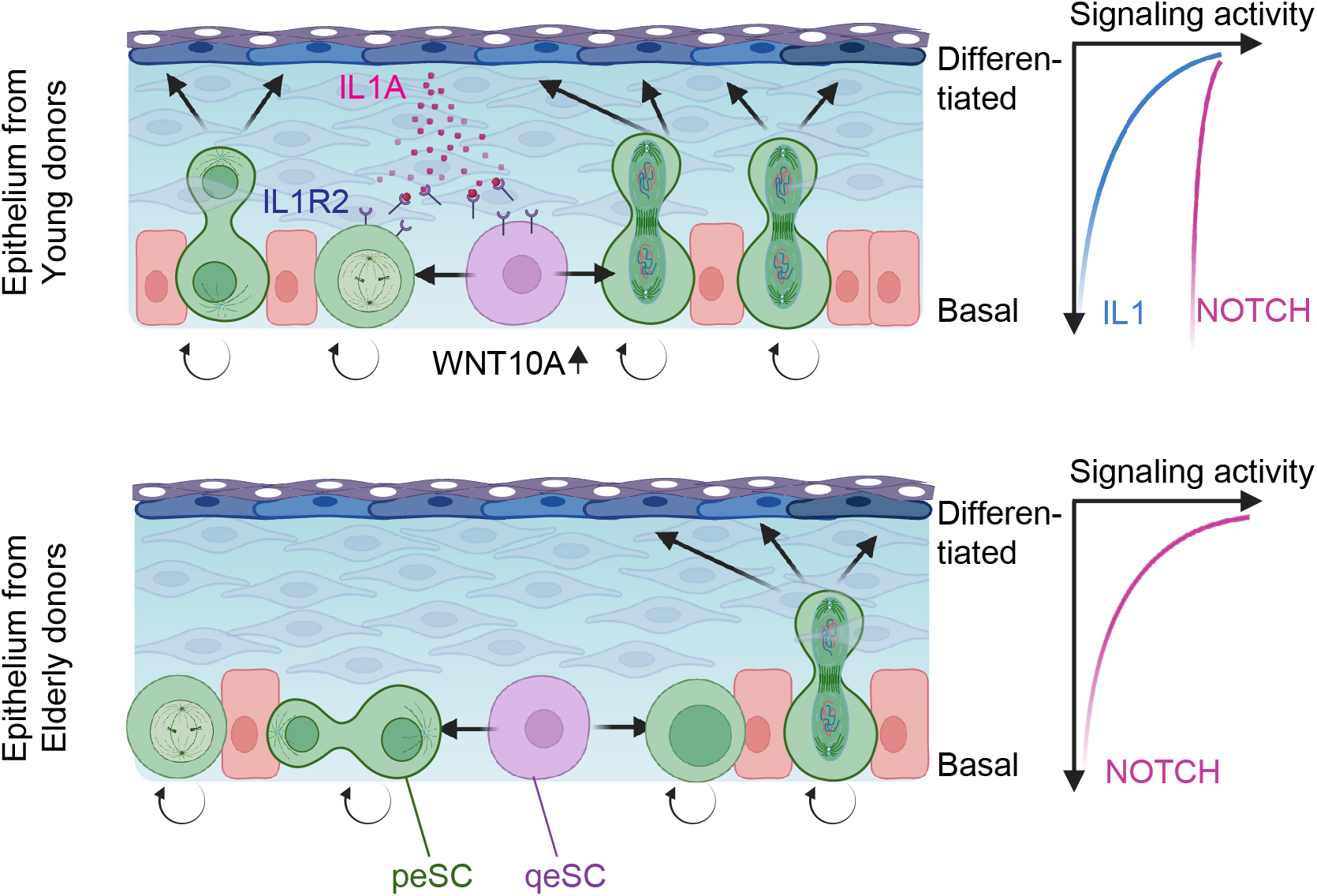
Age-dependent changes in esophageal epithelial homeostasis. Younger and older esophageal epithelia exhibit distinct characteristics within qeSC and peSC populations. In younger tissues, stem cell populations suppress excessive IL1 signaling through expression of decoy receptors while maintaining elevated WNT10A expression and enhanced NOTCH signaling activity. Although both younger and older epithelia contain comparable numbers of stem cells, fewer stem cells actively contribute to differentiation in aged epithelium, resulting in delayed epithelial homeostasis and regeneration.

## Discussion

Esophageal epithelial homeostasis is essential for maintaining epithelial integrity and facilitating tissue repair following damage. Because the esophageal epithelium is continuously exposed to environmental stresses, including smoking and alcohol consumption, preservation of normal lineage differentiation while suppressing abnormal cell growth is critical for maintaining tissue architecture and preventing malignant transformation that can lead to esophageal cancer.

Recent advances in next-generation sequencing and single-cell technologies have enabled the identification of normal esophageal stem cell populations and their associated markers.^1,10,12^ However, age-related differences in stem cell behavior and epithelial homeostasis have not been extensively investigated. In this study, using publicly available single-cell datasets from healthy esophageal epithelium, we analyzed age-dependent differences in stem cell activity and epithelial differentiation dynamics.

Aging is a major factor that impairs epithelial homeostasis and regenerative capacity.^16,17^ In the present study, we sought to identify mechanisms underlying age-associated defects in esophageal epithelial maintenance. Using our VR score–based differentiation simulation, we found that stem cell differentiation dynamics progressively slowed in older age groups (**Fig. 3**). Interestingly, the overall plasticity difference between stem cell and differentiated cell states, represented by the top-to-bottom height in the Waddington landscape, was not substantially altered across age groups, suggesting that intrinsic stem cell plasticity itself is largely preserved during aging. In addition, the overall proportion of stem cell populations did not significantly decline with age. However, differentiation distances became elongated in older tissues, reflected by a less steep slope in the Waddington landscape. Because landscape slope is determined by transcriptional proximity among cell clusters in UMAP space, the increased differentiation distance likely reflects weakened transcriptional coupling between stem and differentiated cell populations in aged tissues. These findings suggest that, although stem cell number and intrinsic plasticity are relatively maintained, fewer stem cells actively contribute to epithelial differentiation in aged tissues. Such reduced functional participation of stem cells may contribute to delayed epithelial turnover and impaired tissue repair during aging.

One of the most prominent age-associated differences identified in this study involved NOTCH signaling activity within the esophageal epithelium (**Fig. 4**). NOTCH signaling is well known to regulate epithelial differentiation,^18^ and loss of NOTCH1 signaling has frequently been observed in aged esophageal tissues.^19^ However, the biological consequences of NOTCH1 loss remain controversial. Previous studies have suggested that NOTCH1 loss may promote clonal competition and suppress esophageal squamous cell carcinoma (ESCC) development,^20,21^ whereas other studies have demonstrated that NOTCH1 inactivation cooperates with TP53 and CDKN2A loss to promote ESCC initiation.^22^ Our analysis showed that NOTCH signaling activity remained broadly preserved in differentiated epithelial cells regardless of age, whereas signaling activity within stem cell populations progressively declined in older tissues. Because NOTCH signaling mediates direct cell–cell communication and epithelial organization, reduced NOTCH activity within stem cell populations may reflect weakened intercellular connectivity and impaired coordination with neighboring cells. Given that many stem-like cells in aged tissues appeared to retain stem cell identity while exhibiting reduced differentiation potential, restoration or enhancement of NOTCH signaling may represent a potential strategy to reactivate quiescent stem cells and improve epithelial homeostasis during aging.

Another notable finding involved IL1- and WNT10A-associated signaling pathways. IL1 family cytokines are known to contribute to gastrointestinal tissue homeostasis and tissue repair.^23^ Because the differentiated esophageal epithelium is continuously exposed to environmental stimuli, including microbiota and tissue injury, IL1-mediated signaling may play important roles in epithelial repair and immune cell recruitment.^23-25^. IL1 signaling has also been implicated in stem cell proliferation and tissue regeneration, including hair follicle regeneration,^26,27^ but excessive pro-inflammatory IL1 signaling is also associated with squamous cell carcinoma and colorectal cancer progression.^28-30^ Similarly, although WNT10A expression has been reported in esophageal cancer cells,^30^ it is also strongly associated with normal developmental stem cell populations, including dental and epidermal stem cells.^31,32^ Interestingly, our analysis demonstrated that *WNT10A* expression was highly enriched within qeSCs specifically in the youngest age group (**Fig. 3B**). Conversely, qeSCs and peSCs in younger tissues preferentially expressed the IL1 decoy receptor *IL1R2*, implying that qeSCs might actively suppress excessive inflammatory IL1 signaling despite the presence of IL1 ligands in the surrounding epithelial environment (**Supplementary Fig. S4B, C**). These findings identify age-dependent alterations in stem cell-associated signaling pathways and provide potential molecular mechanisms underlying impaired epithelial homeostasis during aging.

Despite these interesting findings, this study has several limitations. First, the number of donors included in the analysis was relatively small, which may limit the generalizability of the findings across broader human populations. Additional large-scale datasets will be required for further validation. Second, this study is based primarily on computational analysis of single-cell RNA sequencing datasets and lacks direct experimental validation. Moreover, future studies using functional stem cell assays and in vivo experimental models are necessary to further address the mechanistic roles of NOTCH and IL1 signaling in age-associated esophageal epithelial homeostasis.

In summary, our study provides a comprehensive analysis of age-dependent alterations in normal esophageal epithelial homeostasis at the single-cell level. By integrating lineage trajectory analysis, differentiation simulation, and cell-cell communication prediction, we demonstrate that aging is associated with impaired stem cell differentiation dynamics and altered signaling interactions within the esophageal epithelium. These findings provide insight into the molecular and cellular mechanisms underlying age-associated epithelial dysfunction and offer a foundation for developing strategies to preserve epithelial homeostasis and prevent esophageal disease progression.

## Author contributions

J.J.: Methodology, investigation, software, analysis, data curation, writing, visualization; J.Z.: Investigation; A.P.: Analysis, investigation; S.J.: Investigation; J.-I.P.: Conceptualization, analysis, writing, visualization, supervision, project administration, funding acquisition; K.-P.K.: Conceptualization, methodology, investigation, software, analysis, data curation, writing, visualization, funding acquisition

## Acknowledgments

This work was supported by the National Cancer Institute (CA286761 to K.-P.K. and CA278967, CA278971, and CA296049 to J.-I.P.). Figures were created using BioRender.com.

## Methods

### Esophageal organoid culture

Organoid culture was performed as previously described.^11,22,33,34^ Murine esophageal organoids were cultured in esophageal mouse epithelial organoid medium (EMEOM) for 5 days until characteristic organoid structures, including central keratinized pulp and spheroidal morphology, were established. Organoids were subsequently maintained under different culture conditions. For stemness-enriched conditions, organoids were continuously cultured in EMEOM, whereas WRN-deprived medium was used to induce homeostatic differentiation conditions. After 1 week of culture, organoids were collected for downstream analyses.

### Immunohistochemical analysis

All staining procedures were performed as previously described.^22^ Esophageal organoids (EOs) were harvested by dissociating Matrigel with ice-cold PBS and fixed in 4% paraformaldehyde at room temperature. Following paraffin embedding, organoid sections were mounted onto glass slides. For hematoxylin and eosin (H&E) staining, sections were incubated in hematoxylin for 3–5 min followed by eosin Y for 20–40 s. For immunofluorescence staining, sections were blocked with 3% goat serum in PBS for 30 min at room temperature and incubated overnight at 4°C with primary antibodies against KRT13 (Abcam, 1:250), MKI67 (Abcam, 1:250), SOX2 (Cell Signaling Technology, 1:250), TP63 (Abcam, 1:250), CDH1 (Cell Signaling Technology, 1:400), and β-catenin (Cell Signaling Technology, 1:400). Sections were subsequently incubated with secondary antibodies (1:250) for 1 h at room temperature. Slides were mounted using ProLong Gold Antifade Mountant with DAPI (Invitrogen), and images were acquired using a Zeiss AxioVision fluorescence microscope.

### Preprocessing scRNA-seq and spatial transcriptomic data

Human single-cell RNA-sequencing datasets were obtained from the Gene Expression Omnibus (GEO) and the European Nucleotide Archive (ENA) under accession numbers GSE201153 and PRJEB31843, respectively. Mouse Xenium In Situ spatial transcriptomic data were generated in our previous study (accession number: GSE304762)^14^. Data preprocessing was conducted using Scanpy (version 1.10.4)^35^. Cells expressing fewer than 200 genes and genes detected in fewer than three cells were excluded during quality-control filtering.

### Cell phase Analysis

Cell-cycle phase was assigned using the score_genes_cell_cycle function in Scanpy with S-phase and G2/M-phase gene sets reported by Tirosh et al.^36^ For each cell, S and G2/M scores were calculated relative to matched reference genes, and cells were assigned to S, G2/M, or G1 phase according to the resulting scores. These annotations were subsequently used to assess differences in cell-cycle composition across samples.

### Valley-Ridge score and Waddington-like landscape analysis

Valley-Ridge (VR) scores were calculated as a weighted sum of two components, Valley and Ridge, with weights of 0.9 and 0.1, respectively. VR score computation was performed on a per-sample and per-cluster basis. The Valley component was defined as the median CCAT entropy value, calculated using SCENT^37^, for each sample-cluster combination. To calculate the Ridge component, RNA velocity length was first inverted and scaled between 0 and 1. Cell centrality distance within each cluster was then computed from single-cell PHATE coordinates using the compute_distdeg() function defined by Qin et al.,^38^ with the knn parameter optimized according to cluster size. For each sample-cluster combination, the Ridge component was calculated as the product of the median scaled inverse RNA velocity length and the scaled cell centrality distance. The final VR score was then obtained by combining the Valley and Ridge components. Waddington-like landscapes were visualized using Houdini Indie (SideFX, v20.0.533). VR scores were plotted along the y-axis, and single-cell PHATE coordinates were projected onto the xz plane.

### RNA Velocity–Based Lineage Trajectory Inference

As previously described, RNA velocity, latent time, and PAGA-directed trajectories were analyzed using scVelo^39,40^. Spliced and unspliced Cell Ranger output matrices were merged, and genes with fewer than 20 total counts were excluded. The top 2,000 highly variable genes were selected for downstream analyses. Following size normalization and log transformation, neighborhood moments were computed and RNA velocities were estimated using the dynamical model. Velocity streamlines were visualized on UMAP embeddings.

### Cell differentiation modeling

After generating the 3D models, each Waddington landscape was used for differentiation simulations in Houdini, as previously shown. ^14,15^ Fifty balls were released from the ridge region corresponding to the qeSC or peSC populations. The initial dropping height and physical parameters, including movement speed, were kept identical across all simulations. Balls were colored green while located above the differentiated-cell state and changed to blue upon reaching the differentiated-cell level. Ball movement trajectories were continuously tracked throughout the simulation. The starting point was defined as the frame at which the last of the 50 balls reached the top ridge of the landscape, whereas the endpoint was defined as the frame at which the final ball reached the differentiated-cell state and changed to blue. Representative snapshots were captured together with corresponding frame information.

### Signaling Score Analysis

Pathway activity scores were calculated using the “scanpy.tl.score_genes” function in Scanpy with default parameters as previously decribed. Reference gene sets were obtained from MSigDB database. For NOTCH signaling analysis, genes from the MSigDB ‘HALLMARK_NOTCH_SIGNALING’ (‘APH1A’,‘ARRB1’,‘CCND1’,‘CUL1’,‘DLL1’,‘DTX1’,‘DTX2’,‘DTX4’,‘FBXW11’,‘FZD1’,‘FZD5’,‘F ZD7’,‘HES1’,‘HEYL’,‘JAG1’,‘KAT2A’,‘LFNG’,‘MAML2’,‘NOTCH1’,‘NOTCH2’,‘NOTCH3’,‘PPA RD’,‘PRKCA’,‘PSEN2’,‘PSENEN’,‘RBX1’,‘SAP30’,‘SKP1’,‘ST3GAL6’,‘TCF7L2’,‘WNT2’,‘WN T5A’) gene set were used. For IL1 signaling analysis, genes from the ‘REACTOME_INTERLEUKIN_1_PROCESSING’ (‘CASP1’,‘CTSG’,‘GSDMD’,‘IL18’,‘IL1A’,‘IL1B’,‘NFKB1’,‘NFKB2’,‘RELA’) were used for IL1 signaling score.

### Cell-cell interaction Analysis

Cell–cell interaction inference was performed using the CellChat package. Integrated epithelial, fibroblast, and immune-cell datasets were used to generate CellChat expression matrices. For age-dependent interaction analyses, epithelial cells were isolated and analyzed separately according to donor age group.

## Code availability

The code used to reproduce the analyses described in this manuscript can be accessed via GitHub (https://github.com/jaeilparklab/esophageal_lineages_by_age).

