## Supplementary Information for "Spatiotemporal Transcriptomic Dissection Uncovers Age-Dependent Deceleration of Esophageal Cell Differentiation"

**Supplementary Videos (1–6)**

**Supplementary Figures (1–2)**

### Supplementary Videos

**Supplementary Video 1:** Esophageal epithelial cell differentiation in the 10-15 age group.

**Supplementary Video 2:** Esophageal epithelial cell differentiation in the 30-35 age group.

**Supplementary Video 3:** Esophageal epithelial cell differentiation in the 40-45 age group.

**Supplementary Video 4:** Esophageal epithelial cell differentiation in the 50-55 age group.

**Supplementary Video 5:** Esophageal epithelial cell differentiation in the 60-65 age group.

**Supplementary Video 6:** Esophageal epithelial cell differentiation in the 65-70 age group.

### Supplementary Figure legends

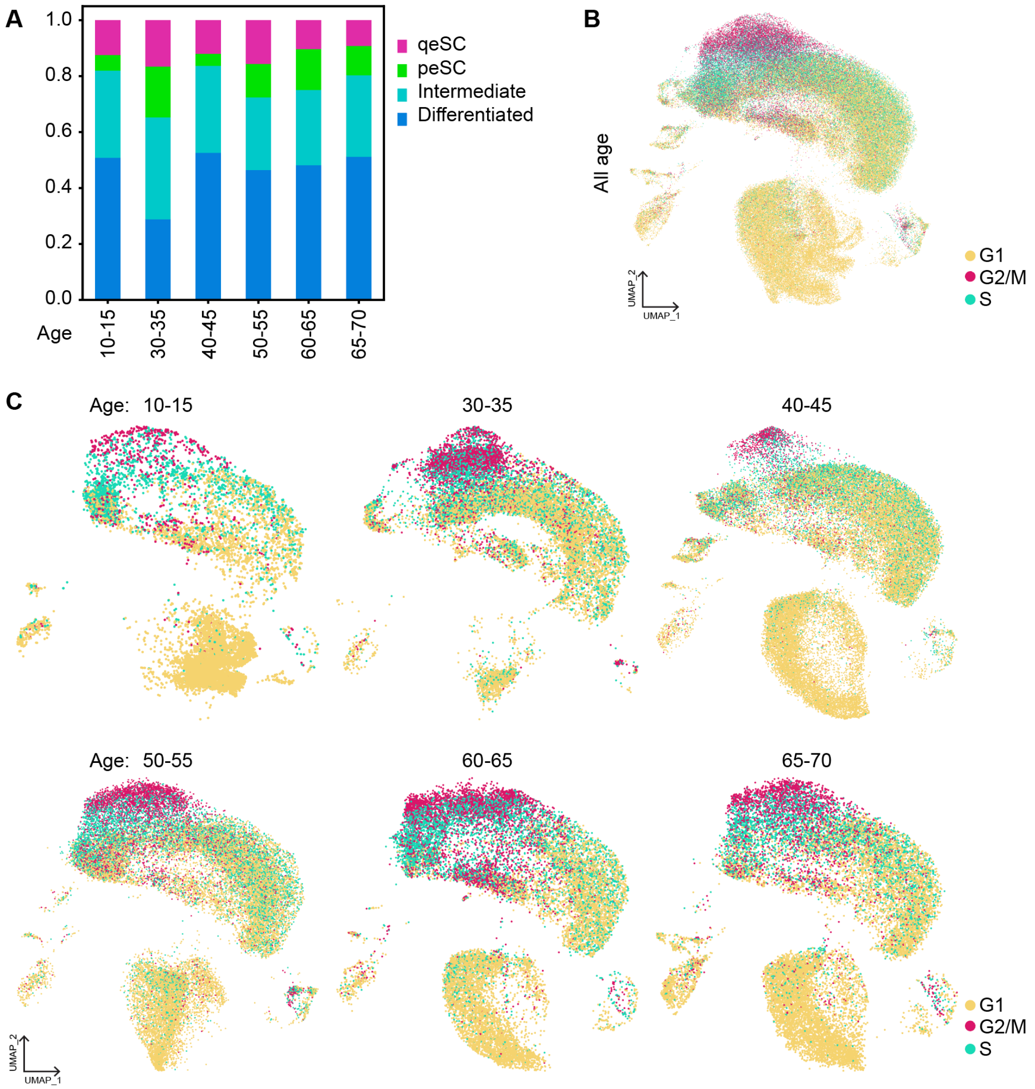


**Figure S1. Age-dependent characteristics of esophageal epithelial cells**

**A.** Proportions of epithelial cell populations across different age groups are displayed as stacked bar plots.

**B.** UMAP visualization showing cell-cycle phases of the entire epithelial cell population.

**C.** UMAPs displaying cell-cycle phase distributions across different age groups.

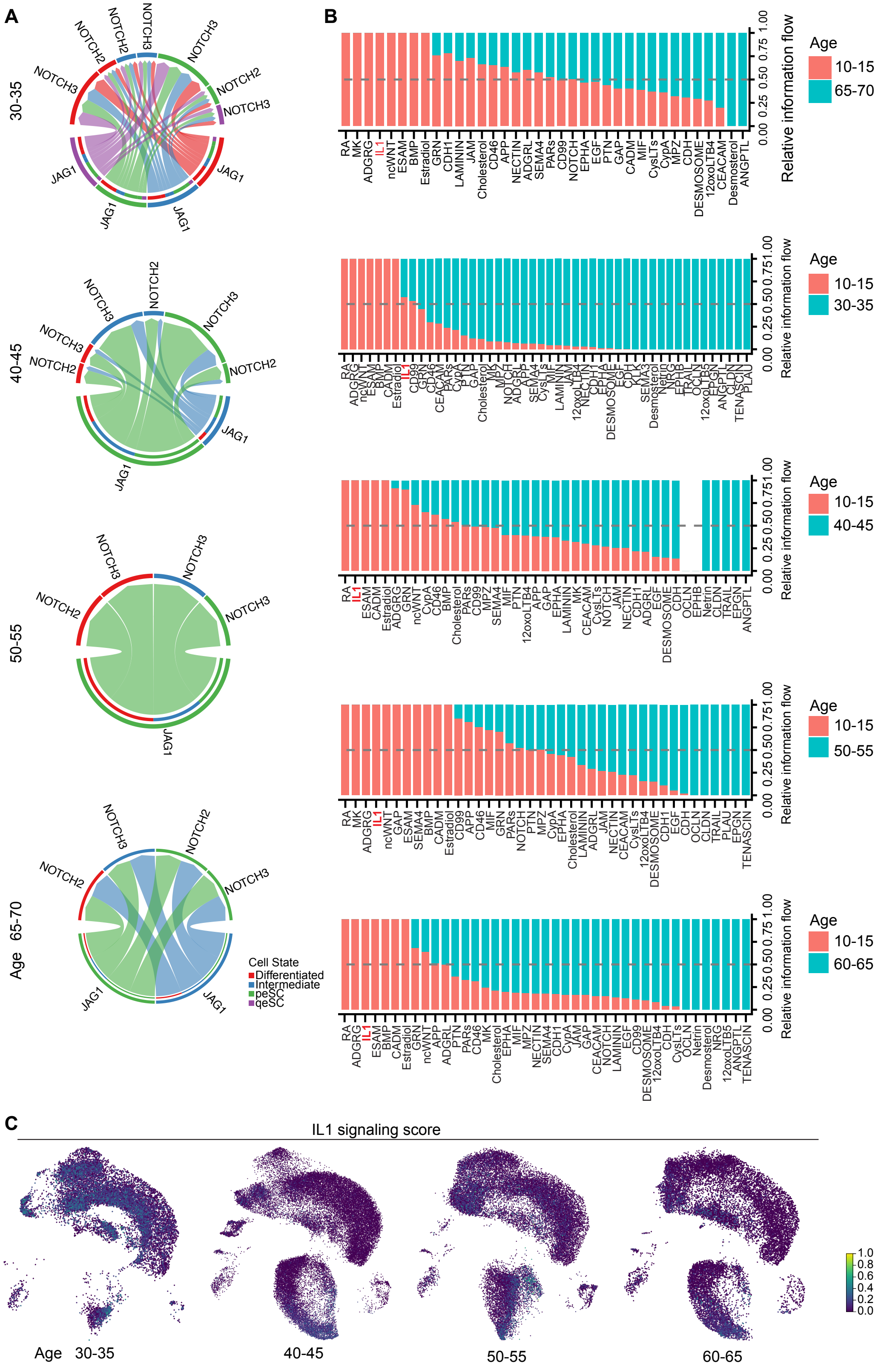


**Figure S2. Age-related signaling patterns in esophageal epithelium**

**A.** NOTCH signaling-mediated cell-cell interaction within the esophageal epithelium across age groups.
**B.** Comparative analysis of significant cell-cell interactions relative to the 10-15 age group.
**C.** UMAPs displaying IL1 signaling activity scores in different age groups.
